# Optimization of swelling response of pH sensitive hydrogels using Box-Behnken design

**DOI:** 10.1101/403857

**Authors:** Fereshteh Shokri, Pouneh Ebrahimi, Soheyla Honary

**Affiliations:** Faculty of Pharmacy, Mazandaran University of Medical Science. Sari, Iran; Department of Chemistry, Gonbad Kavous University, Gonbad, Iran; Department of Pharmacy, Mazandaran University of Medical Science, Sari, Iran

**Author notes:** Corresponding author: Fereshteh Shokri, Faculty of pharmacy, Mazandaran University of Medical Science, Sari, Iran.

**Keywords:** Box-Behnken Design, chitosan, chemometrics, optimization, pH-sensitive hydrogels

## Abstract

Hydrogels are three-dimensional crosslinked hydrophilic polymer networks capable of swelling or de-swelling reversibly in water and retaining a large volume of liquid in swollen state. Hydrogels can be designed with controllable responses to shrink or expand with changes in external environmental conditions. Among stimuli-sensitive hydrogels, the pH-sensitive ones are widely-studied and used. Despite these advances, we currently lack a systematic way to optimize the synthesis protocols. Here we propose to use techniques from chemometrics, namely Box–Behnken statistical screening design to optimize the chemical composition of pH-responsive hydrogels for an improved responsiveness as quantified by swelling index. Polymer blends were prepared by mixing different suitable volume of pvp (poly vinyl pyrrolidone) and chitosan aqueous solution in order to obtain a mixture, glutardialdehyde solution was added as crosslinkers to the chitosan/pvp mixture to form semi-IPN(semi-interpenetrating polymeric hydrogel). The pH-dependent swelling properties have been measured and used to obtain a regression model. We characterized the descriptive and predictive abilities of our model. We found a remarkable correlation (correlation coefficient = 0.954) between the observed responses and the responses predicted by the model. Our results demonstrate that Box-Behnken is an appropriate statistical design that can be successfully used in the development of pH-sensitive hydrogels with a predictable swelling ratio. This technology will remarkably reduce the time and cost that is needed for chemical synthesis of hydrogels with a desired pH-sensitivity.

## 1 Introduction

Hydrogels are polymeric materials that have the ability of swelling and retaining significant fraction of water within its structure, though they will not be dissolved in water. Despite their large water content, hydrogels possess a degree of flexibility like natural tissues. Hydrogels are created from copolymers carrying randomly distributed positively and negatively charged groups [1]. Hydrophilic functional groups attached to the polymeric backbone; increase the hydrogels ability for water absorption while they are resistant to dissolution due to the inter-polymer cross-links. Cross links and interactions between charged groups determine the viscoelastic properties and stickiness of the hydrogels. By changing the composition of the copolymers and the density of cross links, one can modulate various properties of hydrogels including their swelling property. Hence, such systems can be used for multi-phase drug delivery. [2]

Various approaches have been developed for hydrogel synthesis. One of the procedures is polymerization of multifunctional monomers. Multipolymer interpenetrating polymeric hydrogel (IPN), a major class of hydrogels, is made by cross-linking two distinct synthetic and/or natural polymer components, to form a polymeric network. Semi-IPN hydrogel is a type of IPN hydrogels with only one component of the assembly being cross-linked and the other one is a non-cross-linked polymer [3, 4].

Stimuli responsive hydrogels are hydrogels with tunable physical properties. These hydrogels are designed to respond to various physical and chemical stimuli. Physical stimuli include temperature, electric or magnetic field, light, pressure and sound, and the chemical stimuli include pH, solvent composition, ionic strength, and molecular species (Figure 1).

**Figure 1.**
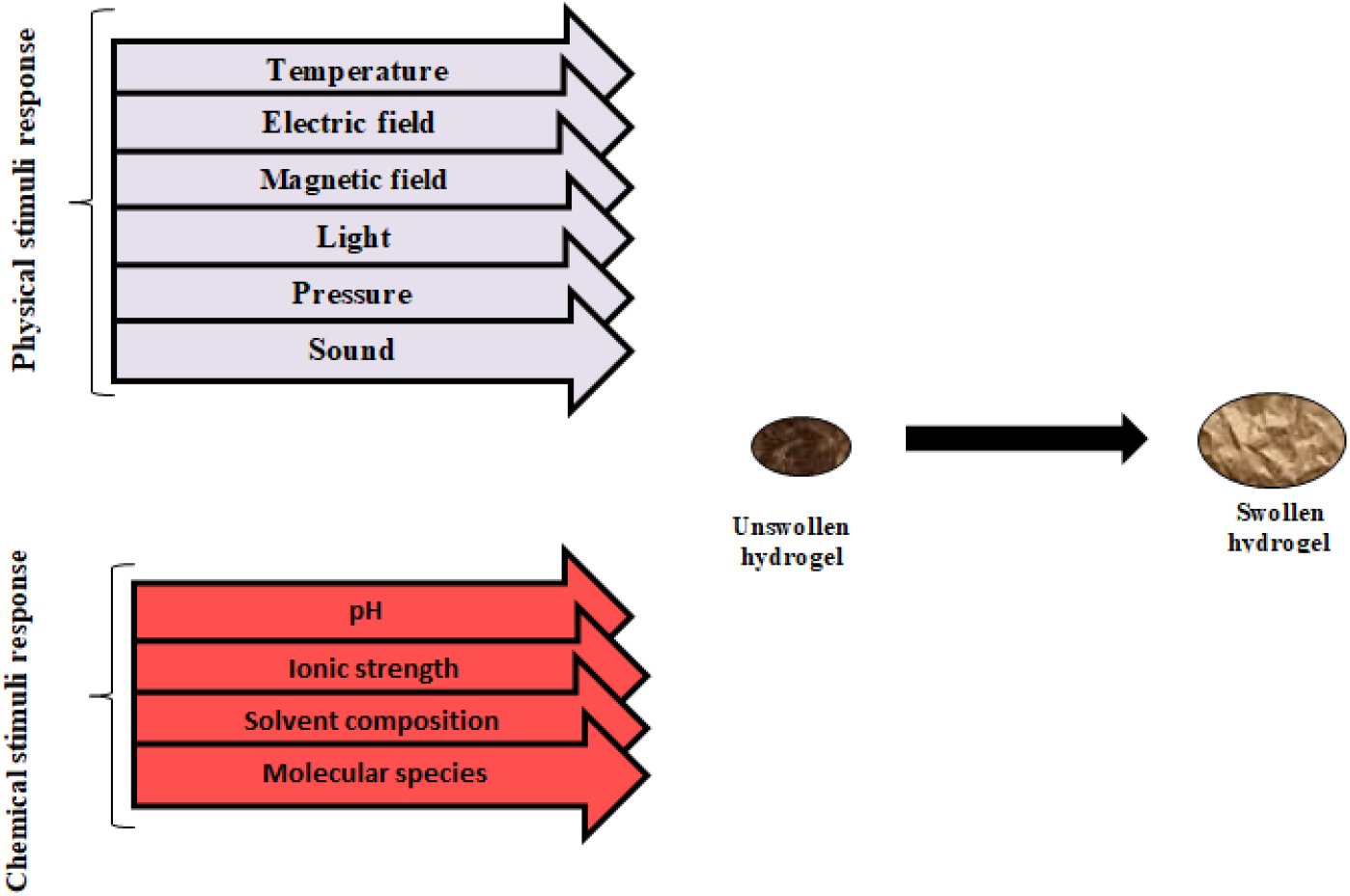
Stimuli induced swelling of hydrogels.

**Figure 2.**
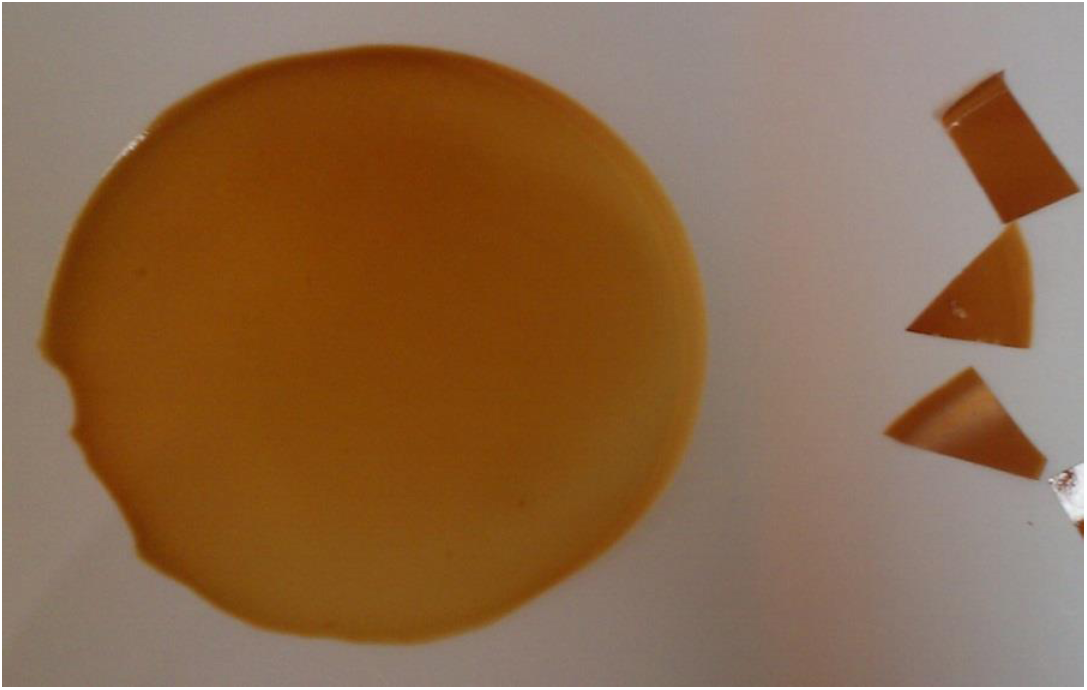
A representative image of the synthesized semi-interpenetrating network polymer (IPN)

Among stimuli-sensitive hydrogels, the pH-sensitive hydrogels have been extensively studied and utilized. Depending on the pH of the surrounding medium, a pH-sensitive hydrogel swells dynamically. The common properties of pH-sensitive hydrogels are the presence of dissociable acidic or basic pendant groups, which can be ionized at certain pH values [5]. A hydrogel with acidic groups swells as the pH level exceeds the acid dissociation constants of the groups. In a similar way, a hydrogel with basic groups swells as the pH level reaches below the basic dissociation constants. pH-sensitive hydrogels have been used widely in biomedical and biotechnological fields [6], especially found useful for controlled drug delivery systems. An important application of pH-sensitive hydrogels is in cancer drug delivery, where acidic pH of the tumor microenvironment (due to local hypoxia, glycolysis and lactate production) induces the release of the drugs from the hydrogel. This approach not only decreases the side effects but also promotes the efficacy of chemotherapy.

Swelling ratio is considered as an important measure for designing and optimizing a hydrogel as a drug delivery system. Response surface methodology (RSM) is one of the methods for the improvement and optimization of drug delivery systems. RSM involves generation of polynomial mathematical relationships and planning of the response over the experimental domain to choose the optimum formulation [7]. Box–Behnken statistical design is a type of RSM design that is an independent, rotatable or nearly rotatable second-ordered design which consists of the central point and the middle points of the edges [8]. Compared with other RSM designs, the Box-Behnken design offers some advantages such as running the variables at only three levels and avoiding treatment combinations that are extreme per se, with some different achieving results. In the present study, we implicated the Box-Behnken experimental design for optimization of pH-sensitive hydrogels based on swelling ratio, wettability and uniformity constant. We focused on chitosan, the result of N-deacetylation of chitin which has an undeniable impact on drug delivery systems, due to its biocompatibility, biodegradability, nontoxicity and low immunogenicity [9]. Chitosan consists of both hydroxyl and amino groups; it can react chemically and hence participated in the salt formation. Blending of chitosan with hydrophilic polymers is likely to increase flux and selectivity of water. These hydrophilic groups are taken account to play significant role for creating a preference water adsorption. Moreover, chitosan hydrogels show how pH-dependent swelling behavior has to do with ionization of amino groups [10]. In previous studies, it has been illustrated that high molecular weight chitosan have high viscosity and low water solubility in neutral pH, thus limiting its applications [11,12]. This higher viscosity may lead to more difficulties in preparing suitable swelling index; fortunately, it can be escaped by using low molecular weight chitosan which is watersoluble in a wide ranging of pH and prepares an acceptable swelling index [13].

## 2 Experimental

### 2.1 Materials and methods

Chitosan with three different molecular weights (150 000, 200 000 and 400 000) was obtained from Fluka Biochimica (Buchs, Switzerland). Poly vinyl pyrrolidone (pvp), gluterdialdehyde, acetic acid, sodium hydroxide, hydrochloric acid, potassium chloride and potassium phosphate monobasic were obtained from Merck, Germany. Double distilled deionized water was used throughout this study.

### 2.2. Preparations of pH-sensitive hydrogels

Chitosan solution (2%, w/v) was prepared in 0.35M acetic acid solution at ambient temperature by stirring for 2hour. pvp solution (4%,w/v) was prepared in deionized water by stirring for 1hour as well [14]. Polymer blend, also, was prepared by mixing different suitable volume of pvp and chitosan aqueous solution in order to obtain a mixture with the following ratios: chitosan/pvp 50/30(v/v), 70/30(v/v), 90/30(v/v(. glutardialdehyde solution was added at various concentration from 0.2, 0.4, 0.6 (v/v) used as crosslinkers to the chitosan–pvp mixture in order to form semi-IPN.hydrogels were cast in a container and then dried by air-drying at 37°. Air-dried hydrogels were prepared by allowing the solvent to be evaporated at 37° for 72 h, then were sliced into pieces and stored.

### 2.3. Preparation of standard Buffer solution

#### 2.3.1: Hydrochloric Acid Buffer

Place 25ml of the potassium chloride solution in a 100-ml volumetric flask, add the 6.5 ml hydrochloric acid solution, then add water to volume for gaining to pH=2

#### 2.3.2: phosphate Buffer

place 25 ml of the monobasic potassium phosphate solution in a 100-ml volumetric flask, add the 2.8 volume of the sodium hydroxide solution, then add water to volume for gaining to pH=6 [15]

### 2.4 Swelling studies

To determine the pH-dependent swelling properties of dried hydrogels, the chitosan/pvp blended hydrogel membrane was sliced into pieces afterwards; pre-weighed dry samples were immersed in solutions with pH values of 2 and 6 at 37°for at pre-determined time intervals (1, 2, 3, 4, 24, 48 hours), swollen hydrogels were removed and then weighed on a sensitive balance.

### 2.5. Swelling kinetics

The amount of water measurement imbibed within the hydrogel is an important criterion for characterizing the hydrogel for biomedical applications, often represented in terms of “swelling ratio” (S) [16]. The swelling ratio is directly proportional to the amount of water imbibed within the hydrogel which influences the diffusional properties of a solute within the hydrogel. Experimentally, the swollen ratio was calculated by the following equation [17]:

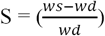

Where *W*s and *W*d are the weight of swollen and dry gel respectively. This procedure was repeated 3 times at all of the time points. Response is the average swelling of the hydrogel in pH=2 divided by average swelling of the hydrogel in pH=6 at the specified times.

### 2.6. The Box-Behnken experimental design and optimization by RSM

The Box-Behnken experimental design with three variables at three level and 15 runs was used for this study. Three factors which had profound influence on the swelling ratio of pH-sensitive hydrogels based on preliminary studies and literature data were selected as the independent variable for this study. The experimental range and levels of the variables were also determined according to the preliminary studies. The independent variables and their levels and the Box-Behnken design matrix are shown in table 1 and 2 respectively. The resulting experimental data were subjected to multiple regression analysis so as to build a regression model for the estimation and prediction of the response. The optimal conditions for the preparation of pH-sensitive hydrogels were obtained by solving the model equation for all possible values of independent variables in the experimental range in this study. The optimal conditions were defined as a condition leading to pH-sensitive hydrogels with largest possible swelling index (low molecular weight chitosan, chitosan/pvp 50/30(v/v) and glutardialdehyde 0.6 (v/v) in this case).

**Table 1.**
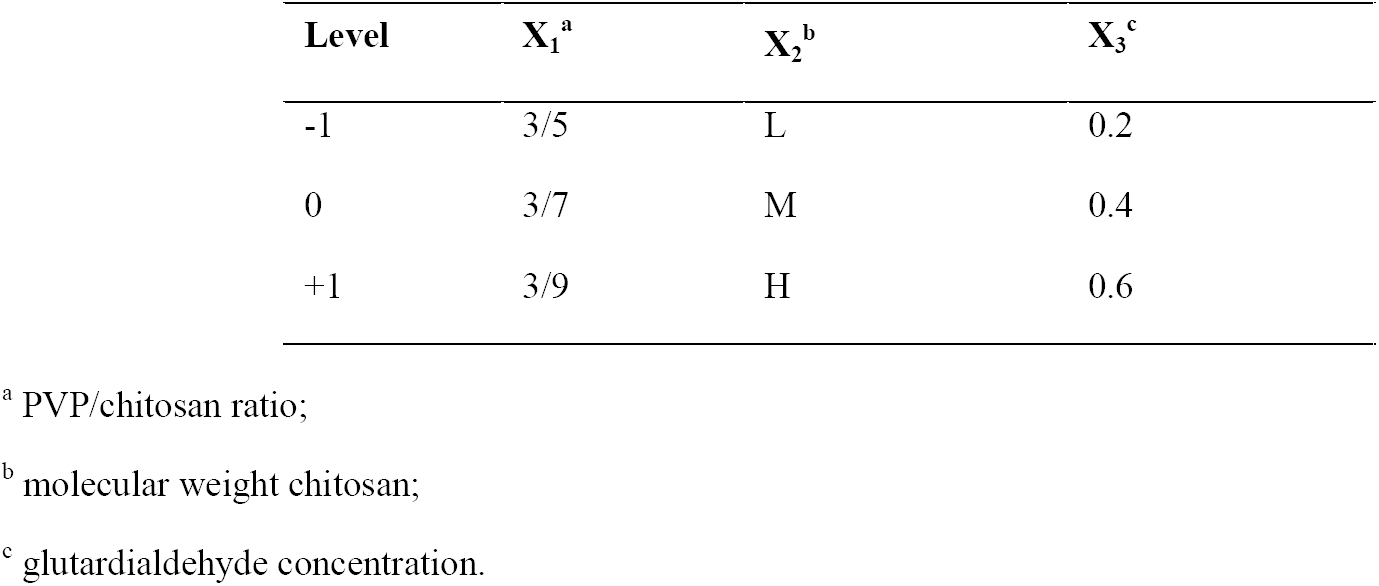
Variables and the selected levels for swelling chitosan/PVP.

**Table 2.** Box-Behnken design for the studied three factors at predetermined time intervals and the experimental results.

| Trial | $X_1^a$ | $X_2^b$ | $X_3^c$ | Swelling ($pH = 2/pH = 6$) | | | | | |
| --- | --- | --- | --- | --- | --- | --- | --- | --- | --- |
|  |  |  |  | 1hour | 2hour | 3hour | 4hour | 24hour | 48hour |
| 1 | + | + | 0 | 0.963 | 1.082 | 0.5910 | 1.040 | 0.799 | 0.930 |
| 2 | + | - | 0 | 1.380 | 0.966 | 1.125 | 1.104 | 1.170 | 1.364 |
| 3 | - | + | 0 | 1.634 | 1.305 | 1.280 | 1.117 | 1.082 | 1.004 |
| 4 | - | - | 0 | 0.910 | 1.184 | 1.095 | 1.064 | 1.154 | 1.470 |
| 5 | + | 0 | + | 1.084 | 1.170 | 1.112 | 1.202 | 1.250 | 1.181 |
| 6 | + | 0 | - | 1.180 | 1.012 | 0.921 | 1.170 | 1.170 | 1.164 |
| 7 | - | 0 | + | 1.070 | 1.338 | 1.017 | 1.128 | 1.225 | 1.281 |
| 8 | - | 0 | - | 1.080 | 1.032 | 1.162 | 1.093 | 1.380 | 1.680 |
| 9 | 0 | + | + | 1.102 | 1.161 | 0.910 | 1.030 | 1.260 | 1.170 |
| 10 | 0 | + | - | 1.120 | 1.410 | 1.412 | 1.618 | 1.166 | 1.119 |
| 11 | 0 | - | + | 1.360 | 1.183 | 0.980 | 1.143 | 0.945 | 1.063 |
| 12 | 0 | - | - | 1.556 | 1.310 | 1.494 | 1.284 | 1.351 | 1.321 |
| 13 | 0 | 0 | 0 | 0.970 | 1.233 | 1.233 | 1.044 | 1.449 | 1.410 |
| 14 | 0 | 0 | 0 | 0.993 | 0.906 | 0.902 | 1.274 | 1.260 | 1.057 |
| 15 | 0 | 0 | 0 | 1.286 | 1.441 | 1.361 | 1.106 | 1.260 | 1.400 |

<sup>a</sup> PVP/chitosan ratio
<sup>b</sup> molecular weight of chitosan
<sup>c</sup> glutardialdehyde concentration

### 2.7. Validation analysis

In order to verify the validity and reliability of the proposed regression model, a number of additional batch validation test in the experimental range of variable in this study were carried out and the observed response were compared to the model predictions.

### 2.8 Statistical analysis

The statistical analysis of data was performed using Microsoft Office 2010 and SPSS for windows, version 16 software packages. The analysis of variance (ANOVA), followed by Fisher’s test, was used for construction and evaluation of the regression model. The standardized effects of the independent variable and their interaction on the dependent variable were evaluated by Student’s t-Test. Value of *P* < 0.05 were considered to denote statistical significant in all cases.

## 3. Results and discussions

### 3.1. Box-Behnken design

Box and Behnken have suggested some designs for a spherical domain. The most particular property of this domain is that each factor takes only three levels. These are based on the construction of balanced incomplete block designs [18]. In this paper, the Box-Behnken design was employed to study the effects of independent variables namely PVP/chitosan ratio (*X*_*1*_), molecular Weight chitosan (*X*_*2*_) and gelutardialdehyde concentration (*X*_*3*_) which are shown in table 1.

Table 2 shows the experimental conditions and the averages of swelling ratio ^*pH*= 2^/_*pH* = 6_ The sequence of experiments was randomized and also the quality of fitted polynomial equation and its adequacy were shown by the coefficient of R^2^ and the value of adjusted-R^2^ of model.

By applying the multiple regressions analysis to the output data the following second-order polynomial equation in a coded form was obtained for explaining the relationship between the response and the independent variables:

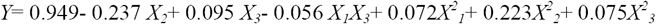

Which *Y* refers to average swelling ratio ^*pH*= 2^/_*pH* = 6_, while *X*_*1*_, *X*_*2*_ and *X*_*3*_ are pvp/chitosan ratio, molecular weight chitosan and gelutardialdehyde concentration respectively. The interaction *X*_*1*_*X*_*3*_ show how the response changes in each single variable, is reflected by the main effect terms (*X*_*1*_, *X*_*2*_ and *X*_*3*_) and quadratic terms 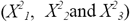

### 3.2. interactions between molecular weight chitosan and swelling ratio

The p-value of molecular weight chitosan, as can be seen in table 4, indicates that it is considered to be statistically significant (<0.05), the β value (−0.237) shows the invers proportionality between molecular weight chitosan and swelling ratio ^*pH*= 2^/_*pH* = 6_. It is interpreted that, an increase in molecular weight chitosan has caused decrease in swelling ratio ^*pH*= 2^/_*pH* = 6_. Findings have proven that low molecular weight chitosan is more appropriate structure than medium and high ones to make pH-sensitive hydrogels for swelling in aqueous media.

### 3.3. Modeling and optimization of swelling ratio

The analysis of variance (ANOVA) was conducted to test the significance and the lack of fit of the quadratic model to the experimental data; the results are presented in table 3. The significance of each model is evident from Fisher’s ratio (F _Model_= 13.46, *P*= 0.001). The F ratio and the corresponding *P* value indicate that the model is able to explain a significant amount of variation in the response. The high value of the determination coefficient (R^2^= 0.910) and the adjusted determination coefficient (adj R^2^ = 0.842) indicate proper fit of the regression model. This shows that up to 91% variability of the response can be explained by the model, leaving only 9% of the total variations in response unexplained. The value of the adjust determination coefficient (adj R^2^) was also high lead to confirm the high important of the model.

**Table 3.** Analysis of variance data for quadratic model.

| Source | SS<sup>a</sup> | df<sup>b</sup> | MS<sup>c</sup> | SE<sup>d</sup> | F<sup>e</sup> | P-value |
| --- | --- | --- | --- | --- | --- | --- |
| Regression | 0.739 | 6 | 0.123 | 0.09569 | 13.459 | 0.001 |
| Residual | 0.073 | 8 | 0.009 |  |  |  |
| Total | 0.813 | 14 |  |  |  |  |

<sup>a</sup> sum of square;
<sup>b</sup> degree of freedom;
<sup>c</sup> means square;
<sup>d</sup> standard error
<sup>e</sup> Fisher's ratio.

**Table 4.** Components of the quadratic regression model and their coefficients.

| Factor | coefficient | standard error | *t* | *p* |
| --- | --- | --- | --- | --- |
| Intercept | 0.949 | 0.055 | 17.178 | 0.000 |
| $X_2^a$ | -0.2370.034 | -6.999 | 0.000 | |
| $X_3^b$ | 0.095 | 0.0342.821 | 0.022 | |
| $X_I^{2c}$ | 0.072 | 0.050 | 1.447 | 0.186 |
| $X_2^{2d}$ | 0.223 | 0.050 | 4.474 | 0.002 |
| $X_3^{2e}$ | 0.075 | 0.050 | 1.502 | 0.171 |
| $X_I X_3^f$ | -0.056 | 0.048 | -1.179 | 0.272 |

<sup>a</sup> mw;<sup>b</sup> g;<sup>c</sup> cpvp<sup>2</sup>;<sup>d</sup> mw<sup>2</sup>;<sup>e</sup> g<sup>2</sup>;<sup>f</sup> cpvpg.

In order to determine the significance of different components of the quadratic model, the standardized effects of the model components were compared using Student’s *t*-test. The results are demonstrated in table 4. According to this table *X*_*2*_ and *X*_*1*_*X*_*3*_ had the minimal effect on the swelling ratio, whereas *X*_*3*_ and *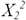* affected the swelling ratio the most as is evident from *t*-value and corresponding *P* value for these terms.

Comparison of the standardized effects of the model components illustrates the most and least significant components of the quadratic model (those components which had maximal or minimal contribution to the generation of response). The *t*-value for each component is a measure of that component’s standardized effect therefore the larger the magnitude of *t*-value and the smaller the corresponding *P*-value for a parameter, the more profound influence on model.

It can be clearly seen in table 4 that although the chitosan molecular weight (*X*_*2*_) showed noticeable effects, with a high value of *t* among all of the components (*t*= -6.999, *P*= 0.000), this parameter had a decreasing effect on the swelling ratio ^*pH*= 2^/_*pH* = 6_due to negative sign. Not only the quadratic of chitosan molecular weight 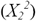had more effects on swelling ration (*t*= 4.474, *P*=0.002) but also glutardialdehyde Concentration (*X*_3_) had this properties, but in second level (*t*=2.821, *P*=0.022). Both of these parameters had maximal effects on swelling index because of positive sign. The glutaraldialdehyde concentration is considered statistically important. The proportionality between crosslinking density and swelling ratio ^*pH*= 2^/_*pH* = 6_can be explained.

An increase in crosslinking density causes growth in level of swelling and pH-sensitivity, by altering the stability of the network, consequence of this trend is a dramatic rise in swelling.

Since the equation is in coded form, the net effect of the independent variables on the swelling ratio could be not inferred from the equation with certainty because at different levels of the variables, the coded values might be negative or positive. This means that based on the variable’s level, the first order and the second order terms for the variable might have synergic or opposite effects on swelling index. Moreover the presence of interaction terms makes the situation even more complicated. So a perfect pattern for the attitude of each independent variable towards increase or decrease in swelling index could not be figured out according to the equation and requires point by point calculation for the entire experimental area.

The parity plot demonstrated in (figure 3), the correlation between the model prediction and the observed responses. The high value of the correlation coefficient (R= 0.954) proved to be remarkable correlation between the observed responses and the responses predicted by the quadratic model. The reliability of the quadratic model was confirmed by the residual plot (figure 4) which shows random distribution of residuals.

**Figure 3.**
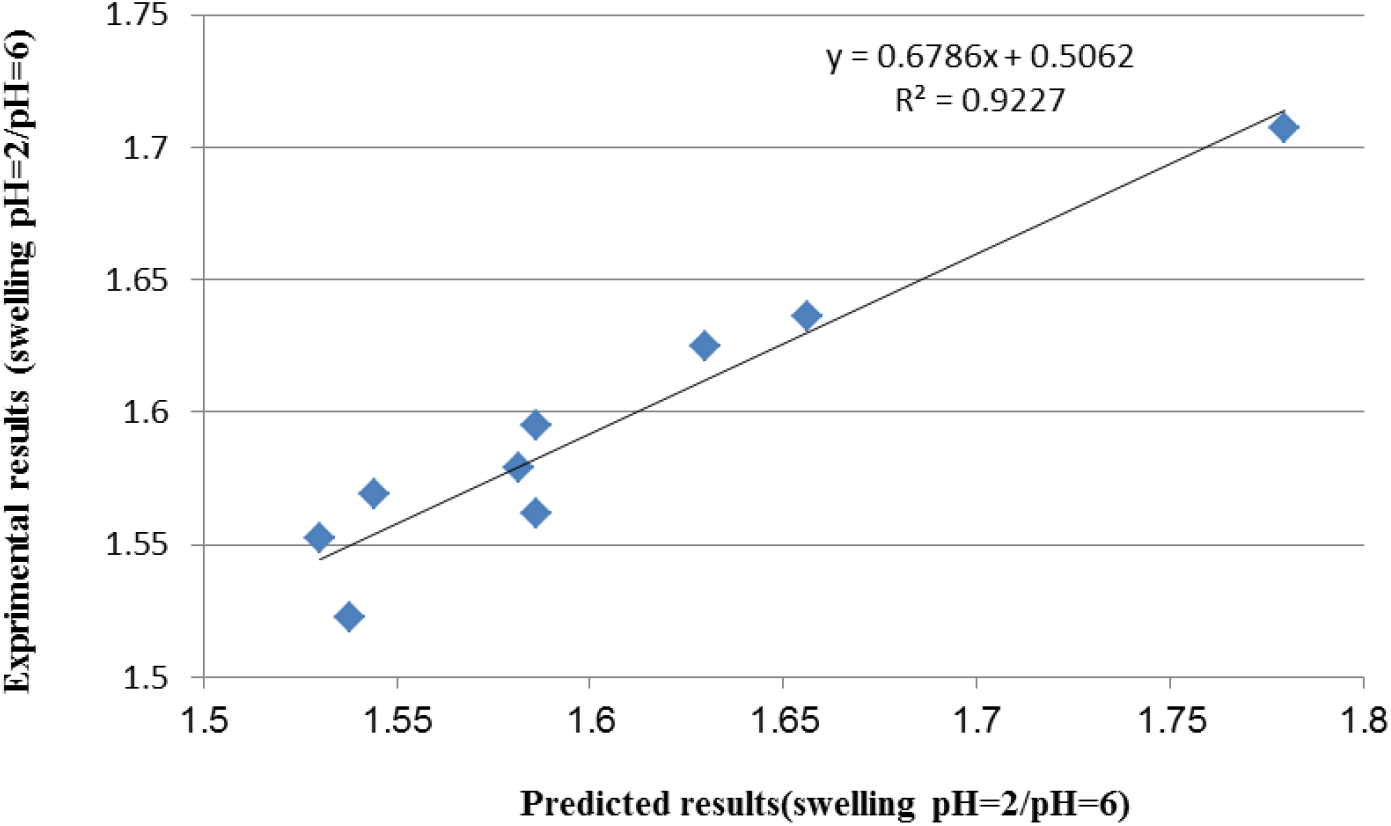
Predicted experimental swelling (^*pH*=2^/_*pH*=6_) for 9 experimental conditions not entered in the modeling.

**Figure 4.**
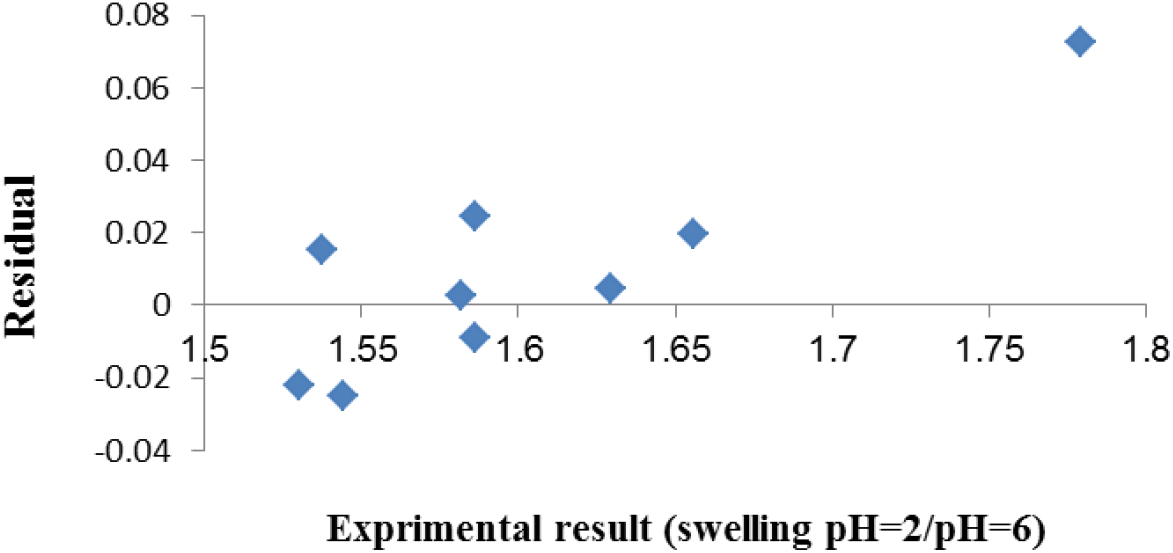
The residual versus plot for the predicted swelling (^*pH*=2^/_*pH*=6_) according to the regression model reported.

According to table 5 the optimum checkpoint formulations were selected to validate the chosen experimental domain and polynomial equations. The optimized check-point formulations were prepared and evaluated for the swelling ratio property.

**Table 5.**
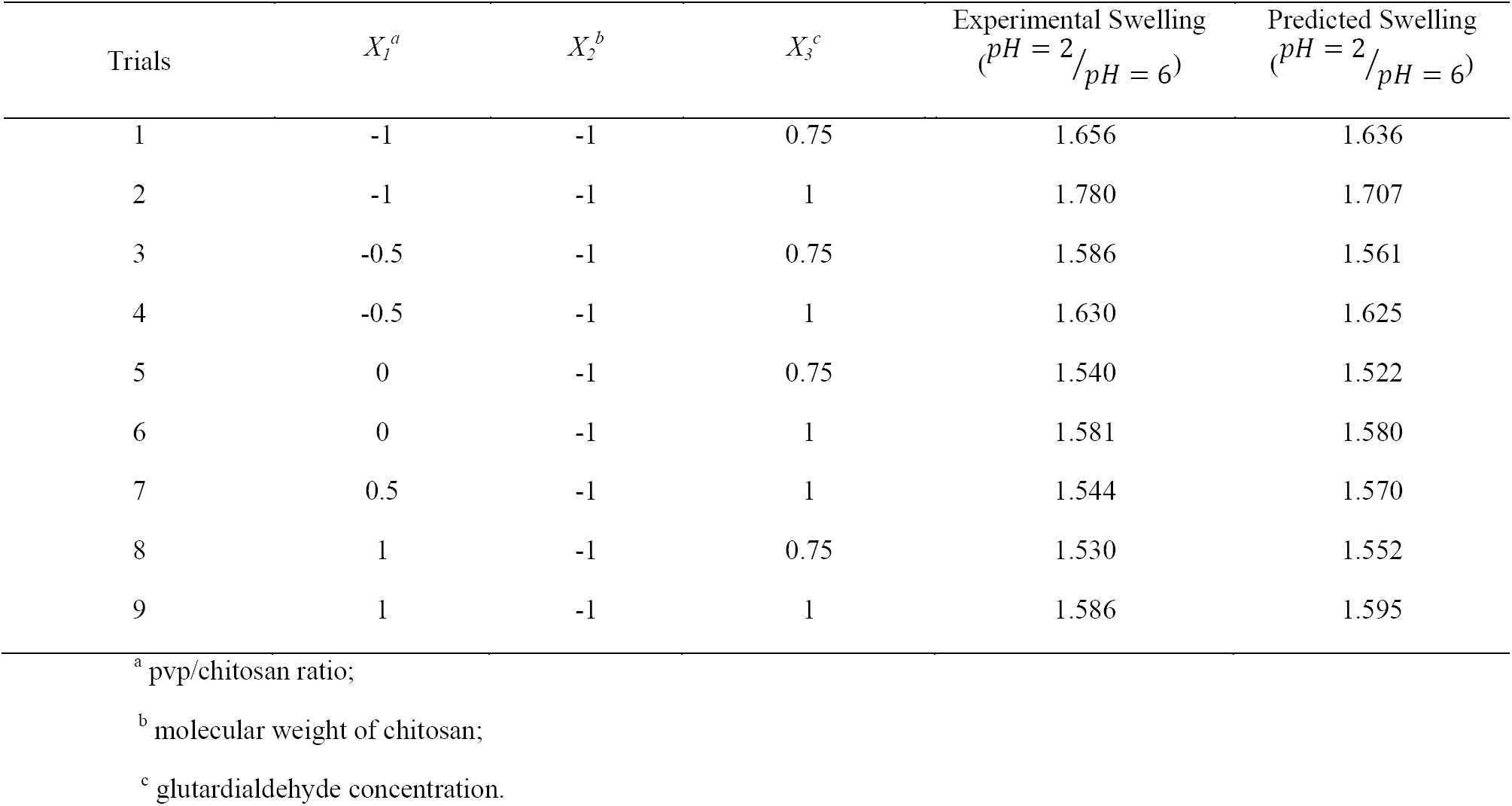
Conditions are used for validating analysis and the observed and predicted responses. The values for independent variables are reported in coded form.

The low molecular weight of chitosan was the most effective factor on swelling ratio.^*pH*=2^/_*pH*=6_ It comes from the fact that swelling process of it is basically completed during the final stage of swelling; erosion process of polymers played an important role in the swelling. Low molecular weight of chitosan is swelled more than high and medium ones. Also, linear regression plots between actual and predicted values of the responses produced (Figure 3). It shows a good agreement between predicted and experimental results for 9 experimental conditions not entered in the modeling. The linear correlation plots drawn between the predicted and experimental values demonstrated high values of R^2^ (0.922) with excellent goodness of fit (p < 0.001). Therefore, the low magnitudes of error and the significant values of R^2^ in the present investigation prove the high prognostic ability of the Box-Behnken in the same rate.

Figure 4 shows the residuals versus of the predicted response. This plot of residuals versus predicted response is essentially used to spot possible heteroscedasticity (non-constant variance across the range of the predicted values), as well as influential observations (possible outliers).

To verify the adequacy of the model to fit the experimental data the compatibility of residuals should be checked out. If the residuals are aligned in the plot then the normality assumption is satisfied. On the other hand, under ideal circumstances, the plots at the top row would not show any systematic structure in the residuals. The histogram would have a symmetric and normal probability plot in a straight line. Figure 4 reveals no apparent problems with the normality of the residuals. To add more information, any systematic structures are not expected in this plot, which would otherwise suggest some transformation (of the response variable or the predictor) or the summation of higher-order (e.g., quadratic) terms in the initial model.

This finding points out notable predictions of maximum responses along with constant variance and adequacy of the quadratic models. Therefore, it can be concluded that the empirical model is adequate to describe average swelling ratio ^*pH*=2^/_*pH*=6_

### 3.4. Influence of pH on swelling

As expected, lower of swelling is encompassed by pH=6 when compared with the pH=2. This conclusion was had to do with the pH-dependent swelling ratio ^*pH*=2^/_*pH*=6_. The swelling ratio ^*pH* =2^/ _*pH* =6_ of high and medium molecular weight chitosan was slower than that of low molecular weight chitosan in pH=2 and pH=6 medium.

## 4 Conclusions

To put it in a nutshell, the pH-sensitive hydrogels of chitosan are successfully prepared with chitosan, pvp and gelutardialdehyde as crosslinker and optimized using a three-factor, three-level Box–Behnken design. The quantitative effect of these factors at different levels on the swelling rate could be predicted by using polynomial equations. The quadratic RSM studied for the swelling ratio helped in comprehending the interaction effects between the combination and ratio of the two polymers. The predicted values were linearly in good agreement with the experimental values of the response variables obtained from the optimized formulations. In this study, the most important factors investigated during the formulation of hydrogel-based swelling index are their mechanical strength and response-time in a physiological environment. Validation of the optimization technique represented the dependability of the model. Moreover, low molecular weight chitosan showed excellent biological activities compared to high and medium ones. We envision that the results of this study will be used to synthesize drug delivery vehicles, and in particular semi-IPNs pH sensitive hydrogels for anti-cancer drug delivery.

## Compliance with ethical standards Conflict of interest

The authors declare that they have no conflict of interest.

